# Triose phosphate utilization stress during photosynthesis addressed with dynamic assimilation measurements

**DOI:** 10.1101/2022.11.27.518113

**Authors:** Alan M. McClain, Thomas D. Sharkey

## Abstract

Oscillations in CO_2_ assimilation rate and associated fluorescence parameters have been observed alongside the triose phosphate utilization (TPU) limitation of photosynthesis for nearly 50 years. However, the mechanics of these oscillations are poorly understood. Here we utilize the recently developed Dynamic Assimilation Techniques (DAT) for measuring the rate of CO_2_ assimilation to increase our understanding of what physiological condition is required to cause oscillations. We found that TPU limiting conditions alone were insufficient, and that plants must enter TPU limitation quickly to cause oscillations. We found that ramps of CO_2_ caused oscillations proportional in strength to the speed of the ramp, and that ramps induce oscillations with worse outcomes than oscillations induced by step change of CO_2_ concentration. An initial overshoot is caused due to a temporary excess of available phosphate. During the overshoot, the plant out-performs steady state TPU and ribulose 1,5-bisphosphate regeneration limitations of photosynthesis but cannot exceed the rubisco limitation. We performed additional optical measurements which support the role of photosystem I reduction and oscillations in availability of NADP^+^ and ATP in supporting oscillations.

**Highlight:** Rapid CO_2_ changes cause more oscillations of photosynthetic rate than a step change in CO_2_ or slowly changing CO_2_. Photosystem I acceptor side limitations may play a role.

## 1. INTRODUCTION

The triose phosphate utilization (TPU) limit on photosynthetic rate can appear when plants are capable of producing phosphorylated Calvin-Benson cycle intermediates faster than these intermediates can be dephosphorylated and converted into end-products (Sharkey, 1985a). When photosynthesis is TPU-limited, inorganic phosphate is not released from the organic phosphate pool fast enough to sustain maximum throughput of both the ATP synthase and the Calvin-Benson cycle, photosynthesis must be downregulated to balance the two. This regulation imposes a cap on the rate of CO_2_ fixation at the rate of end-product synthesis. Plants are not typically TPU-limited under ambient conditions (Sage & Sharkey, 1987; Ellsworth *et al*., 2015). TPU limitation is easiest seen by elevating the rate of photosynthesis through increased light level and CO_2_ partial pressure or decreased O_2_ partial pressure (Sharkey *et al*., 1986b) such that the photosynthetic rate is increased by 10 or 20% relative to ambient conditions (Yang *et al*., 2016). It is more likely to be observed when photosynthesis is measured at a lower temperature than growth conditions (Stitt, 1986; Sage & Sharkey, 1987; Labate & Leegood, 1988), due to the high temperature sensitivity of end product synthesis (Stitt & Grosse, 1988; Leegood & Edwards, 1996), which exceeds the temperature sensitivity of the other biochemical processes in photosynthesis (Cen & Sage, 2005; Sage & Kubien, 2007). The occurrence of TPU limitation depends greatly on the species and the acclimation of the plant. For example, plants grown at low temperature are often resistant to TPU limitation because they develop additional sucrose-phosphate synthase activity (Cornic & Louason, 1980; Guy *et al*., 1992; Holaday *et al*., 1992). Expressing *Zea mays* sucrose-phosphate synthase in tomato significantly reduced the temperature at which TPU was evident (Laporte *et al*., 2001).

TPU limitation is associated with a variety of regulatory processes. TPU-limited plants exhibit reduced rubisco activation state in as little as one min after imposing TPU conditions (Sharkey *et al*., 1986a). Rubisco deactivation can restore the balance between the capacities to fix carbon and convert the fixed carbon to end-products. TPU-limited plants also develop an elevated transthylakoid proton-motive force *(PMF)* and an associated increase in energy-dependent quenching (Takizawa *et al*., 2008; Kiirats *et al*., 2009). This increase is probably associated with declining phosphate concentration in the stroma (Sharkey & Vanderveer, 1989) driving up the ΔG_ATP_ of the stromal ATPase reaction. One consequence of this regulatory arrangement is the reduction of *φ_PSII_* as [CO_2_] increases (Sharkey *et al*., 1988; Stitt & Grosse, 1988). The requirement for electron transport is determined by the rate of photosynthesis and photorespiration. Increasing [CO_2_] reduces the rate of photorespiration under TPU-limited conditions but the increased availability of electron transport capacity cannot be used resulting in a decline in *φ_PSII_*. The decline in *φ_PSII_* balances the rate of electron transport with the reduced requirements for electrons because of the reduced rate of photorespiration.

In TPU-limited photosynthesis, photosynthetic rate is defined by regulatory features. To detect TPU limitation in gas exchange data, it is easiest to determine the presence of regulatory mechanisms, such as the increase in non-photochemical quenching or the decline in *φ_PSII_* upon increasing CO_2_ (McClain & Sharkey, 2019), or the CO_2_- or O_2_-insensitivity of the CO_2_ assimilation rate *(A)*, which demonstrates that *A* is not defined by rubisco properties and which is characteristic of TPU limitation (Sharkey, 1985b). These regulatory mechanisms can have different time constants. For example, Sharkey et al. (1986) observed depletions of ATP and RuBP and reductions in ATP/ADP ratio and rubisco activation state 1 min after imposing TPU limitation. However, after 18 min, RuBP was higher than before imposing TPU conditions and the ATP/ADP ratio, and rubisco activation recovered partially. Thus, as different regulatory mechanisms are induced upon imposition of TPU limitation, there can be transients in the specific process setting the rate of photosynthesis, for example the availability of RuBP at one time versus the activation of rubisco at another time.

One consequence of these transients is oscillations in *A*, which have been observed under TPU limitation (Ogawa, 1982; Walker *et al*., 1983; Sivak & Walker, 1986, 1987).Oscillations are commonly seen when the environmental conditions are rapidly changed to elevate the photosynthetic rate, such as a step change in CO_2_ partial pressure or light availability, or a reduction in O_2_ partial pressure (Harris *et al*., 1983) to increase carbon fixation by reducing photorespiration. Oscillations are visible in both carbon assimilation and fluorescence parameters, demonstrating parallel changes in the Calvin-Benson cycle and electron transport (Walker *et al*., 1983; Peterson *et al*., 1988; Stitt & Grosse, 1988). There have been a few models proposed to explain oscillations in photosynthetic rate. In general biological oscillatory models, oscillations are typically caused by a delay in a feedback component of a multiple component system, leading to overshooting of steady-state before inhibition can be achieved. One theory is that there is a delay in activation of sucrose synthesis after a photosynthetic increase (Laisk & Walker, 1986). Another theory is that the delay originates from fructose-2,6-bisphosphate inhibiting fructose-1,6-bisphosphatase (Stitt *et al*., 1984; Laisk & Eichelmann, 1989; Laisk *et al*.,1989).

The use of ramps of CO_2_ to induce oscillations should allow us to study the phenomenology of oscillations with high-speed measurements of *A* and *Ci* (Stinziano *et al*., 2017). However, the 100 ppm/min limit on ramp speed with the Rapid *A/C_i_* Response (RACiR, Stinziano *et al*., 2019) technique combined with inaccurate *C_i_* measurements, especially at the beginning and end of curves, limited this approach. Dynamic assimilation techniques (DAT, Saathoff & Welles, 2021) is a natural evolution of RACiR that features a greater range of ramp rates and better accuracy, especially at the start and end of the ramp. Dynamic calculations of assimilation, which include an accumulation term to account for changes in the concentration of CO_2_ in the chamber that is disregarded in steady-state equations, also make measurements of assimilation possible following sharp changes in [CO_2_]. With DAT, we can now use advanced ramps and step changes in [CO_2_] to clarify the mechanism by which TPU limitation causes oscillations, and how exactly the assimilation rate can surpass the steady-state limit.

## 2 MATERIALS AND METHODS

### 2.1. Plant materials and growth

*Nicotiana benthamiana* seeds were germinated in 2 l pots of potting media consisting of 70% peat moss, 21% perlite, and 9% vermiculite (Suremix; Michigan Grower Products Inc., Galesburg, MI, USA) in a greenhouse from June to August. This greenhouse was located at 42°43’N, 84°28’W, East Lansing, Michigan. Typical daylight light levels were between 300-700 μmol m^-2^ s^-1^, and the daytime temperature was controlled to 27°C during the day. Plants were watered with half-strength Hoagland’s solution (Hoagland & Arnon, 1938) as needed as seedlings and then daily as adults. Plants were used for experiments from 6-7 weeks of age, and the uppermost fully expanded leaves were used for gas exchange.

### 2.2 Dynamic Assimilation Techniques

Dynamic measurements of gas exchange were made in a LI-COR 6800 with a LI-COR 6800 12A 3 cm x 3 cm clear top chamber (LI-COR Biosciences Inc., Lincoln, NE, USA). Plants were acclimated at experimental conditions until steady state with 1000 μmol m^-2^ s^-1^ photosynthetically active radiation and an air flow rate of 800 μmol s^-1^. Dynamic calibrations and range match were performed as recommended in the LI-COR 6800 version 2.0 manual (https://licor.app.boxenterprise.net/s/kt6wwzmnvnlu4vc004pzp9u7cv9bvzj8 pp. 9-66 – 9-109). For experiments presented here, CO_2_ was ramped at rates of 100 to 500 ppm/min (approx. 10 to 50 Pa CO_2_/min. Typical atmospheric pressure was 98 kPa).

### 2.3 Combined optical measurements with gas exchange

A LI-COR 6800 12A 3 cm x 3 cm clear top chamber (LI-COR Biosciences Inc., Lincoln, NE, USA) was connected to a scattering optic with an array of LEDs behind it (Hall *et al*., 2013; Lantz *et al*., 2019). The LI6800 12A backplate was replaced with a 3D-printed plate containing an optical and an infrared detector. The LED array contained actinic red and blue lights producing up to 2500 μmol m^-2^ s^-1^ with a ratio of 90% red (630 nm) and 10% blue (480 nm) light at 1000 μmol m^-2^ s^-1^. The saturation flash provided approx 15,000 μmol m^-2^ s^-1^. Electrochromic shift measurements were made with a 520 nm LED, with 505 nm used to correct for changes in zeaxanthin. The PSII operating efficiency (ϕ_II_) (Baker, 2008) was assessed by chlorophyll fluorescence using 520 nm as the excitation light. Measurements of PSI absorbance were made at 820 nm. While the absorbance at 820 nm may include other signals, such as reduced pheophytin or ferredoxin, these species are in low proportion and change more slowly than P700^+^ and should not significantly affect the kinetics (Christof & Ulrich, 1994). Measurements of PSI were taken according to Kanazawa *et al*. (2017) and measurements of ECS were taken according to Takizawa *et al*. (2007).

To take optical measurements along with the dynamic ramp of CO_2_, plants were first acclimated at 400 ppm CO_2_ and 1000 μmol m^-2^ s^-1^ light until steady state was achieved. CO_2_ was then abruptly lowered to 50 ppm CO_2_ at the reference IRGA, and the plant acclimated at this CO_2_ level for 60 s. Afterwards, CO_2_ was ramped at a rate of 400 ppm/min (approx. 40 Pa/min) (or other rates as indicated) until 1500 ppm CO_2_ in the reference IRGA was recorded. (Because CO_2_ assimilation is a function of partial pressure, assimilation rates are reported as a function of partial pressure. However, the LI-COR 6800 mixes gases in terms of mole fraction, so in explaining experimental design CO_2_ levels are given in ppm). Typical atmospheric pressure at the site of experimentation was 98 kPa and was measured at the time of experimentation for exact calculations. A list of times from 20 to 140 s in 10 s intervals was randomized, and individual ramps were performed sequentially for each interval, allowing assimilation to return to steady state at ambient CO_2_ before beginning the next ramp. At the chosen time, PSI and PSII activity, as well as the dark interval relaxation kinetics of the electrochromic shift (ECS), were measured.

## 3 RESULTS

### 3.1 Oscillations are intensified when induced through ramps rather than CO_2_ step changes

The photosynthetic rate oscillated when the CO_2_ partial pressure was increased sufficiently to cause TPU limitation. When CO_2_ was ramped at rates of 100 to 500 ppm min^-1^, oscillations were more pronounced than when CO_2_ was increased in a step change (Figure 1). The higher amplitude/lower damping oscillations caused by a ramp up of CO_2_ resulted in a lower integral of *A* compared to an abrupt increase (Table 1). Oscillations induced by ramping CO_2_ resulted in on average a 20% loss of total assimilation compared to the steady-state over the course of the ramp, significantly less at p=0.95. Oscillations induced by a step change of CO_2_ performed comparably to the steady state assimilation value at the same CO_2_ level, no significant difference at p=0.95. We fitted a line through the middle of the oscillations. This midline trended down when oscillations were induced by a ramp of CO_2_ but trended up when CO_2_ was changed abruptly.

**Figure 1.**
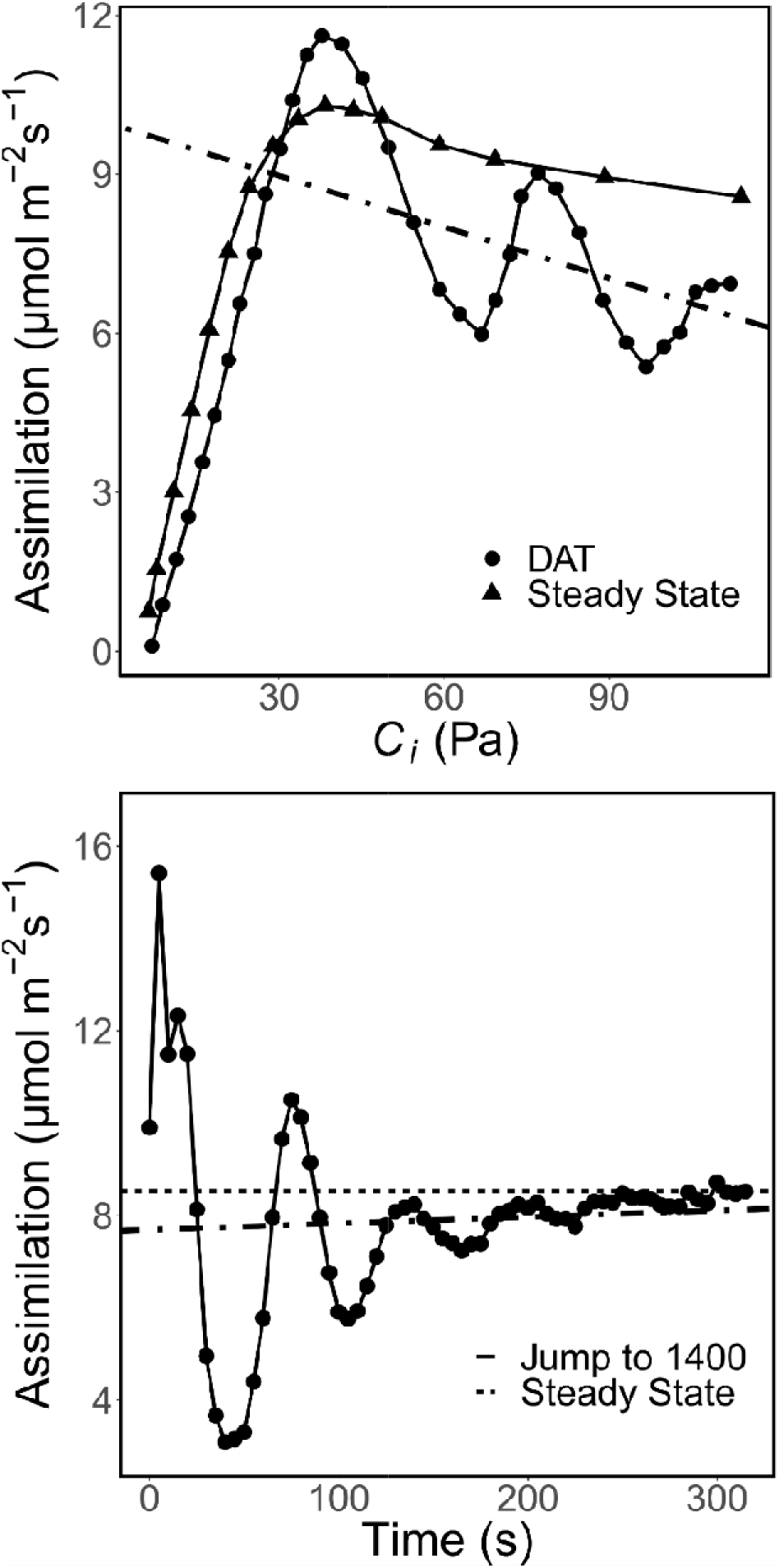
Oscillations induced by elevated CO_2_ compared to the steady state. Top: Full ramp of CO_2_ from 50 ppm to 1500 ppm at a rate of 400 ppm/min compared to a steady-state *A/C_i_* curve (each measurement took 1 to 3 min for the steady-state method). Bottom: Oscillations induced by step-change of CO_2_ from 50 ppm to 1400 ppm compared to steady-state assimilation rate at 1400 ppm CO_2_. For both, a linear model is fit to the oscillating data to show the midline of oscillations.

**Table 1.** A comparison of the total integrated assimilation during oscillations relative to the steady-state assimilation.

| Type | Mean difference<br>(%) | Difference s.d. | 95% CI |
| --- | --- | --- | --- |
| Ramp | -20.0 | 2.6 | -25.1 to -15.0 |
| Step change | -2.2 | 3.9 | -9.9 to 5.5 |

### 3.2 Oscillations are induced specifically by entering TPU limitation

Oscillations were observed only when plants entered TPU limitation (Figure 2). Plants were acclimated at 400 ppm CO_2_ and either 25°C (Figure 2 top and middle) or to prevent the occurrence of TPU limitation, 35°C (Figure 2, bottom). Plants were then exposed to a ramp through a range of CO_2_ values, starting at either 1500 ppm (high to low, labeled backwards, Figure 2, top) or 50 ppm (low to high, labeled “Forwards”, Figure 2 middle and bottom) at a rate of 400 ppm min^-1^. When measured at growth temperature and a ramp from low to high CO_2_, oscillations were observed beginning at a *C_i_* of approx. 30 Pa. When ramped high to low, the plant did not exhibit oscillations at all. When ramped at a higher temperature to prevent TPU limitation, the plant did not exhibit oscillations. Therefore, the oscillations are caused specifically by entering TPU limitation, rather than any of the individual environmental conditions the plant experiences. Leaving TPU conditions does not result in oscillations.

**Figure 2.**
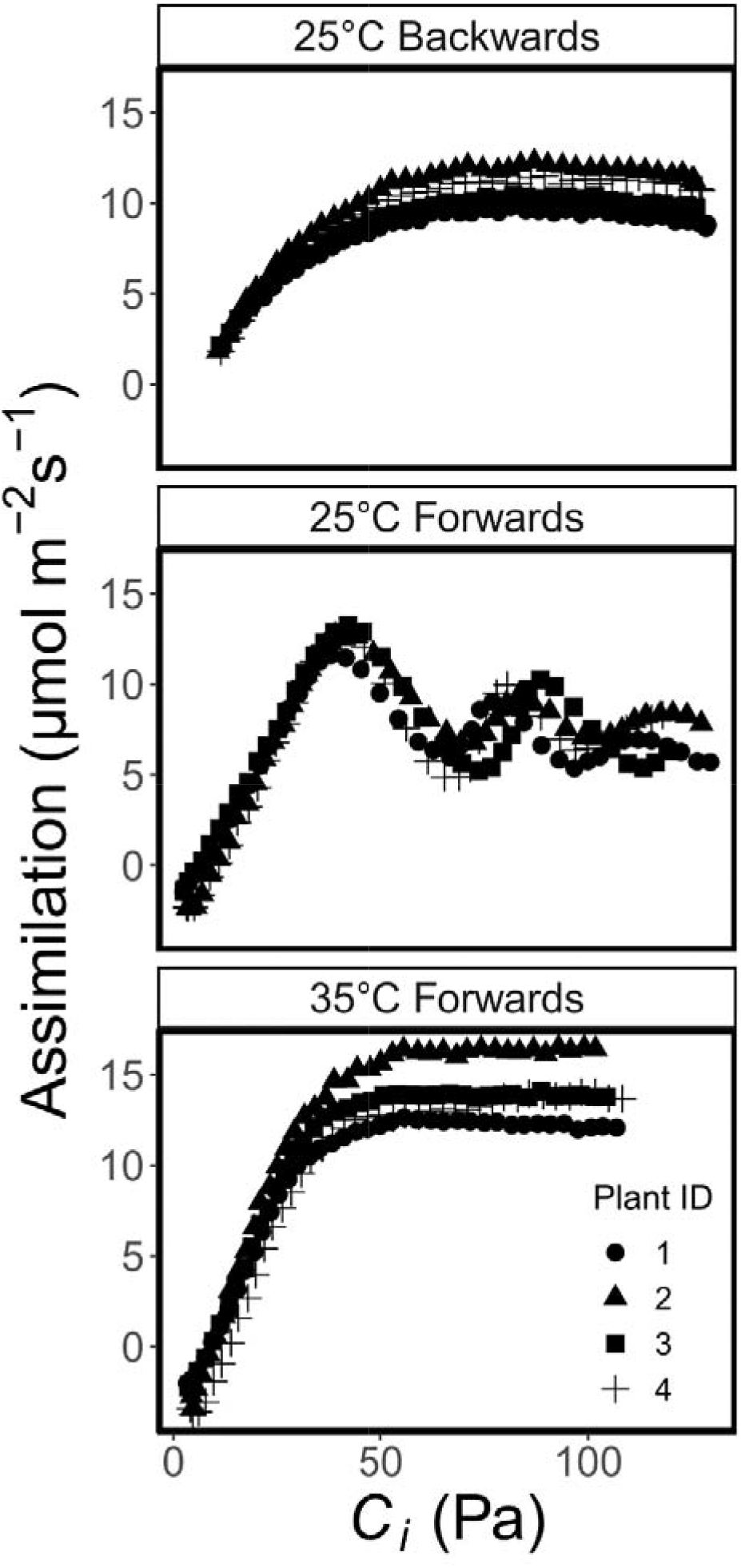
Assimilation measured using dynamic assimilation technique ramps of CO_2_ in three styles. Top: Reference CO_2_ is ramped from 1500 ppm to 50 ppm at 25°C. Middle: Reference CO_2_ is ramped from 50 ppm to 1500 ppm at 25°C. Bottom: Reference CO_2_ is ramped from 50 ppm to 1500 ppm at 35°C. For all curves, CO_2_ is ramped at a rate of 400 ppm/min. Assimilation and *C_i_* are logged every 5 seconds. Different symbols indicate replicate leaves.

### 3.3 Oscillations are intensified when the ramp rate is increased

Plants were acclimated at ambient conditions, then after a 1-min delay at 50 ppm CO_2_, were ramped at a variable rate to 1500 ppm CO_2_ (Figure 3). Sustained oscillations are not observed at a ramp rate of only 100 ppm CO_2_/min but an initial overshoot was seen. The height of the peak of the first oscillation increased with ramp rate. The initial peak value of *A* was greater than the steady state rate except at 100 ppm/min. The initial peak value increased with ramp speed, however, there was a corresponding increase in the depth of the following trough in assimilation. In Figure 3 the assimilation rates are plotted versus *C_i_* but there is also a time element given the variation in the rate of CO_2_ ramp. Supplemental Figure S1 shows the same assimilation rates as in Figure 3 but as a function of time (we put time on a log scale for convenience). Figure S2 shows that the peak assimilation rate decreases with time to reach said peak.

**Figure 3.**
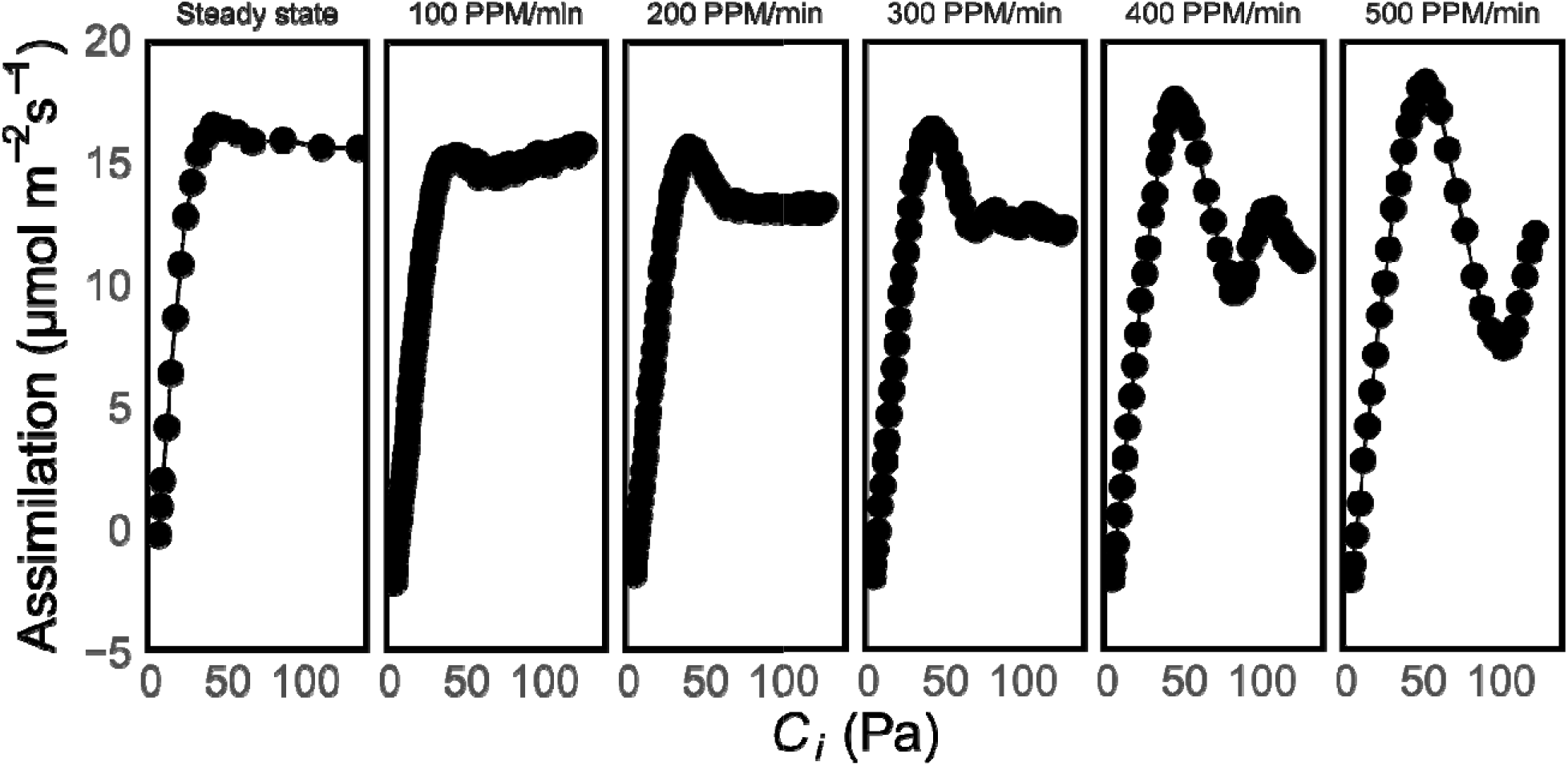
An example set of DAT ramps at various ramp rates, compared against the steady-state *A/C_i_* curve. Reference CO_2_ is ramped from 50 to 1500 ppm at rates of 100 to 500 ppm/min at 25°C. For the steady-state *A/C_i_*, 18 points were collected over a range of reference CO_2_ values from 50 to 1500 ppm over a period of 2.9 – 14.5 min. The amplitude of the oscillations increases in proportion to the ramp rate.

### 3.4 Oscillations are intensified when TPU is enhanced through low temperature

Plants were acclimated until steady state at 20°C at 400 ppm CO_2_, then held at 50 ppm CO_2_ for one min before ramping from 50 to 1500 ppm CO_2_ at a variable rate (Figure 4). The peak amplitudes compared to the steady state were higher relative to those found at room temperature (p<0.1 by Welch’s t test) when ramped at 300-500 ppm/min. However, the absolute peak height is no different (p>0.1 by t test) from the absolute peak height of ambient temperature ramps at 300-500 ppm/min, despite being lower (p<0.1 by t test) at 100-200 ppm/min, as well as in the steady state. Additionally, the ramp rate required to achieve overshooting was lower, 200 ppm min^-1^ rather than 400. These two components combine to increase the oscillation amplitude through the connecting factor of TPU capacity, even though they affect TPU limitation in different ways.

**Figure 4.**
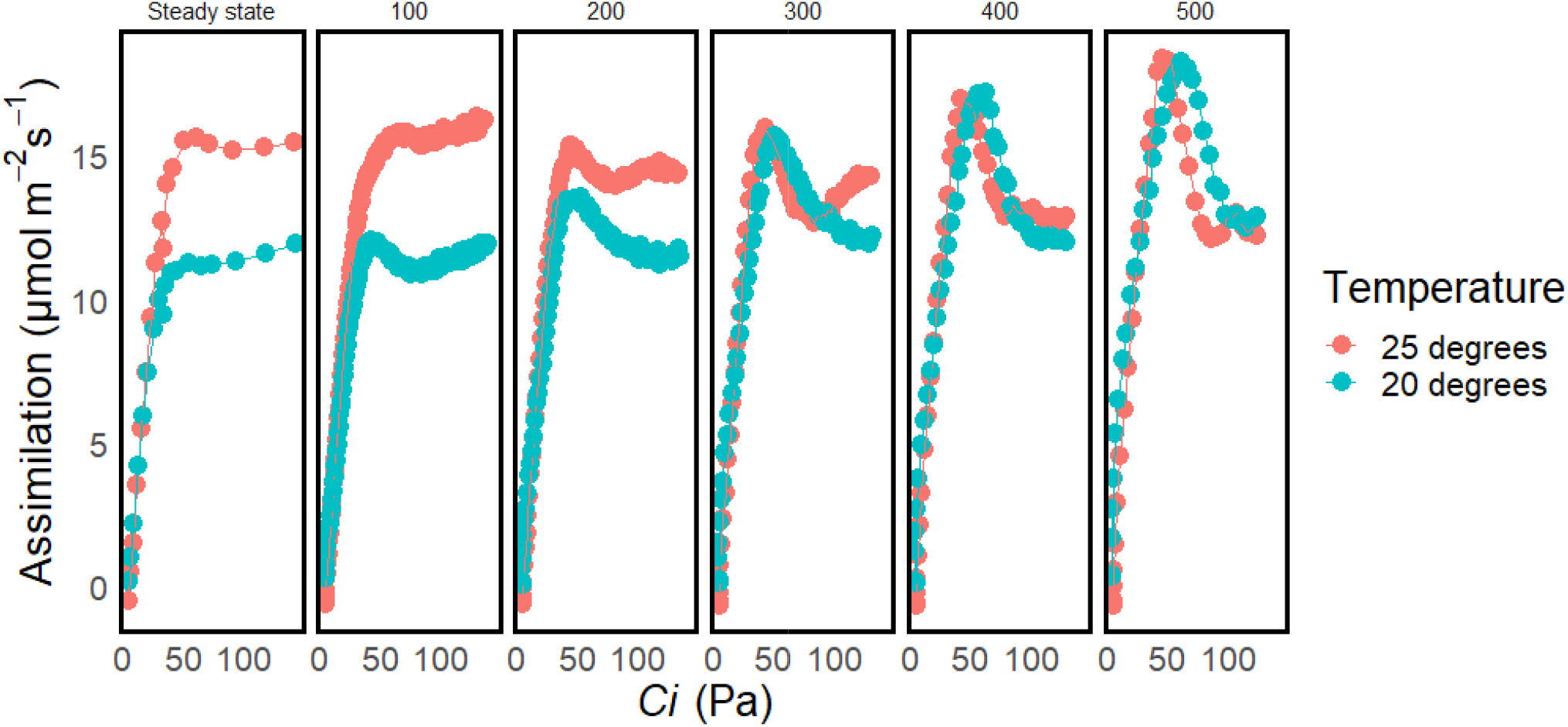
A set of DAT ramps at reduced temperature compared to ramps performed at ambient temperature. Reference CO_2_ is ramped from 50 to 1500 ppm at rates of 100 to 500 ppm/min, compared to an 18-point steady-state *A/C_i_*, all at 20°C. The amplitude of the induced oscillations increases with ramp rate, and is also greater than the amplitude of oscillations at 25°C.

### 3.5 Overshooting dynamically exceeds both TPU and the electron transport limitation of photosynthesis

The oscillations caused by the CO_2_ ramp were plotted with limitations modeled from curve fitting (Gregory *et al*., 2021) for data measured at discreet CO_2_ concentrations. Peak dynamic *A* often exceeded the steady-state TPU limitation during a ramp of CO_2_ (Figure 5). At higher ramp rates, peak dynamic *A* also exceeded the RuBP regeneration limitation of photosynthesis. However, at no point did the overshoots exceed the rubisco limitation of photosynthesis.

**Figure 5.**
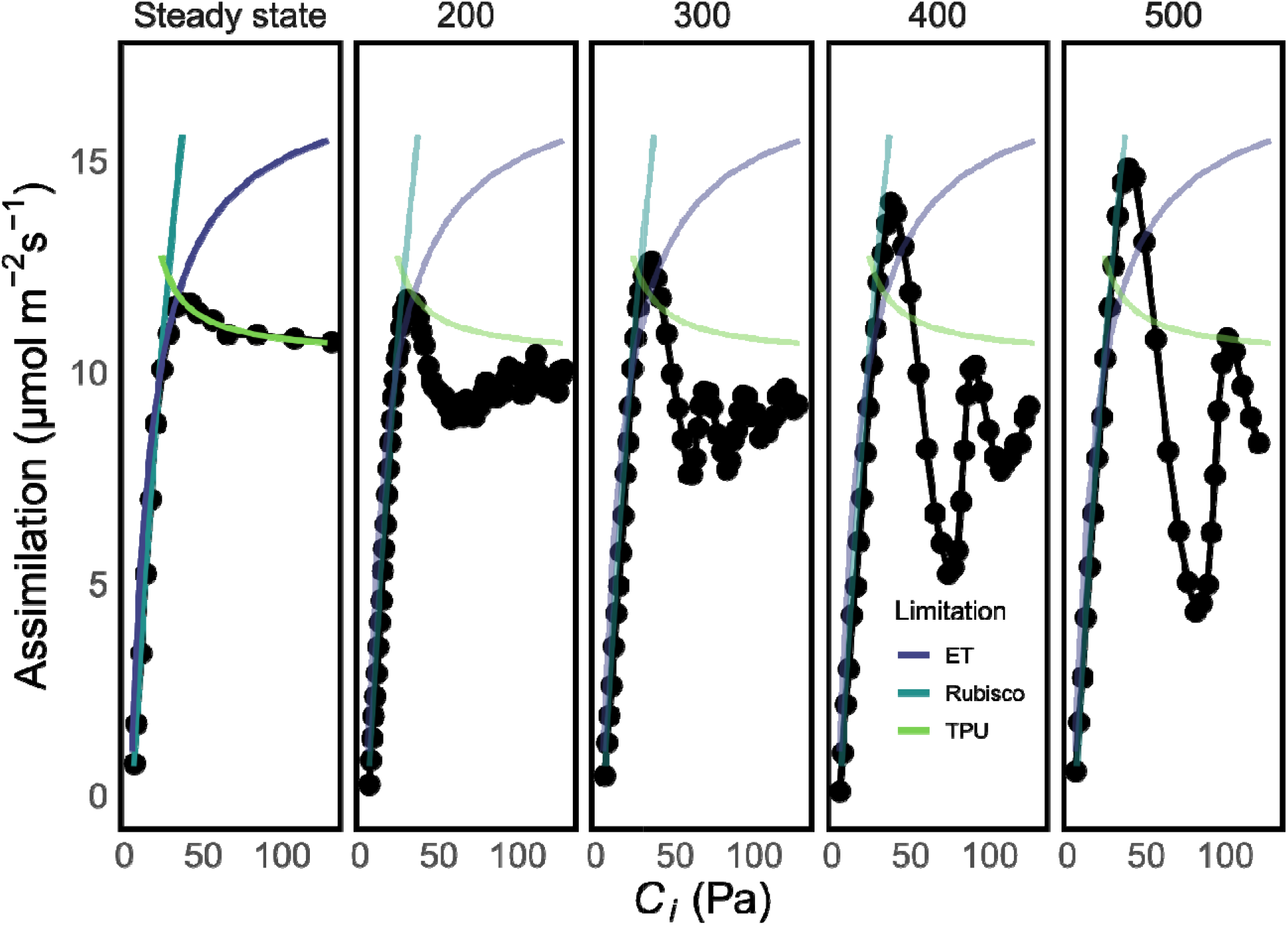
Comparison of oscillations versus fitting parameters from the steady-state *A/C_i_*. Oscillations are induced by ramping from 50 ppm to 1500 ppm at rates varying from 200 ppm/min to 500 ppm/min. Oscillations can easily surpass TPU limitation, and at higher ramp rates can surpass the RuBP regeneration limitation but cannot surpass the rubisco limitation. At the highest ramp rates, the entire overshoot closely matches the rubisco limitation.

### 3.6 PSI reduction was involved in oscillations during CO_2_ ramps

Plants were ramped from 50 to 1500 ppm CO_2_ in a special chamber adapted to house an LED array for measuring electrochromic shift and PSI oxidation in combination with PSII fluorescence based on components of the IdeaspeQ (Hall *et al*., 2013) (Figure 6). Assimilation and *ϕ_II_* were correlated, as previously seen. However, PSI oxidation remained constant throughout the ramp until the first trough, at which point PSI oxidation fell (PSI became reduced). This suggests that the availability of NADP^+^ to accept electrons from PSI became limited.

**Figure 6.**
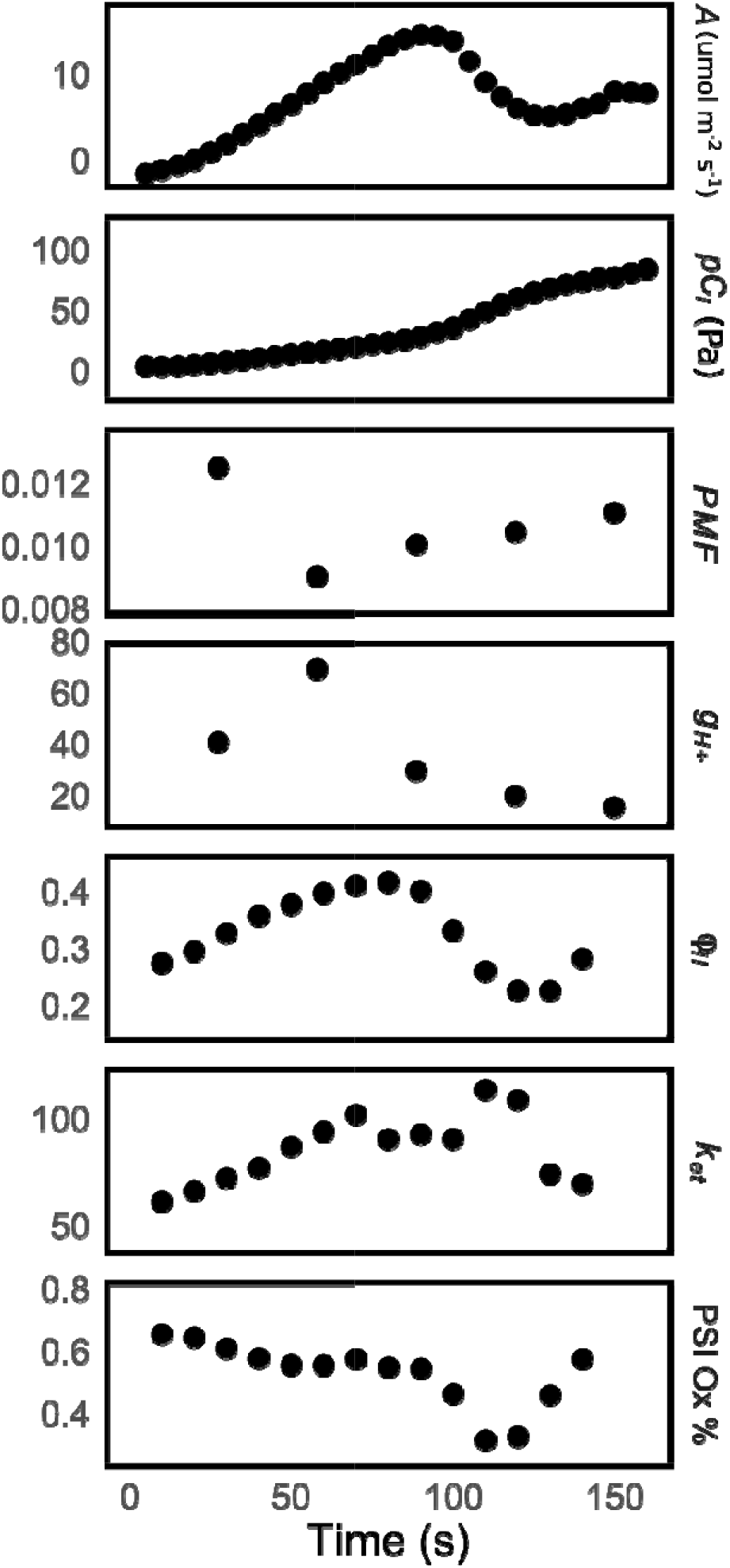
Combination of optical measurements with DAT. Oscillations are induced by ramping from 50 ppm to 1500 ppm at 400 ppm/min. *φ_II_* and PSI oxidation state are calculated from saturation flashes. *PMF, g_H+_*, and *ΔA820_t_* are calculated from dark interval kinetics. *g_H+_, φ_II_* and PSI oxidation state correspond with assimilation, but *PMF* responds in the reverse.

## 4 DISCUSSION

Historically, most of the photosynthetic oscillations research has been performed using sudden shifts in environmental conditions to induce oscillations. The use of ramps of varying speeds helps describe the phenomenology of oscillations to a greater degree, with some implications on the mechanisms of oscillations. The amplitude of oscillations resulting from ramps are greater and the oscillations damp more slowly than oscillations resulting from step changes (Table 2). Oscillations produced by step changes of CO_2_ tend towards the steady state assimilation value. Oscillations produced by ramps, however, tend towards a different midline that diverges from the steady-state assimilation rate. We propose that this is due to the continuous change of the requirements for photosynthetic regulation, which is the damping force of these oscillations. The amplitude of the oscillations is also affected by the rate of the ramp. If the ramp is too slow, overshooting can still occur but not oscillations. In this situation, a simple damped harmonic oscillator model cannot describe the behavior, as overshooting is not seen in an overdamped or critically damped model, and an underdamped model cannot account for the following trough.

**Table 2.**
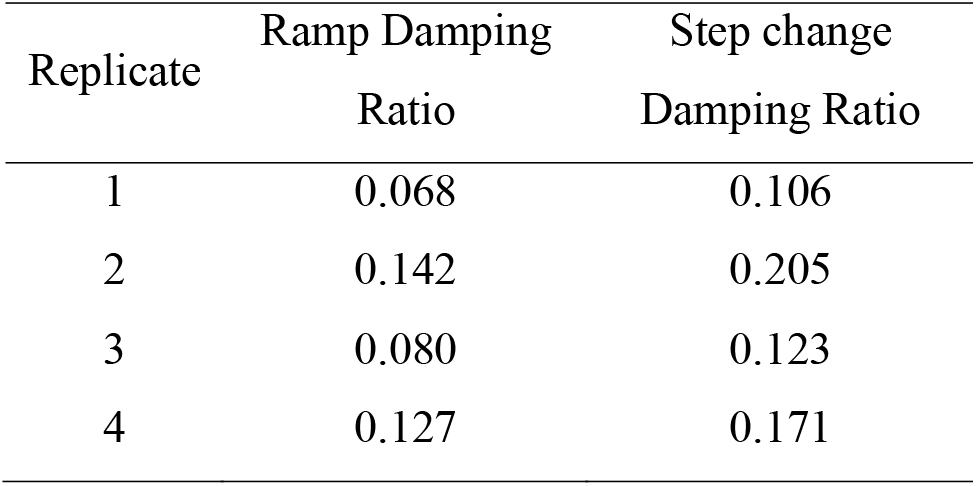
A comparison of the harmonic oscillator damping constants from a set of four plants, with each being tested in both oscillations induced by CO_2_ ramp and a step change in CO_2_. The damping constants were estimated by logarithmic descent of peak height. The mean difference is not 0 at p=0.95 using a two-sided paired t-test (95% CI 0.0291 – 0.065).

The use of ramps also allows us to compare the oscillations to the photosynthetic limitations fit from steady-state behavior. The peak exceeds the RuBP regeneration limitation and the TPU limitation, both of which are functions of metabolite pools. For short periods of time, metabolites such as RuBP can be used faster than they are produced, depleting the pool and adding instability to the system. However, the rubisco limitation is not a function of metabolite pools, it is believed to represent the kinetics of RuBP-saturated rubisco and be unaffected by changes in RuBP pool size (Farquhar, 1979; Sharkey, 2022). It is therefore unsurprising that oscillations did not exceed the rubisco-limited portion of the curve.

Similar transient peaks in *A* above the steady-state rate of RuBP regeneration were induced by short periods of CO_2_-free air (Ruuska *et al*., 1998). Short dark periods can also allow photosynthesis in subsequent light periods to exceed its steady state rate for short periods (Stitt 1986). On this basis, we propose that the overshooting achieved during oscillations results from the transient reduction in pools of metabolites which would otherwise be consumed at a steady-rate, allowing photosynthesis to temporarily exceed the steady state rate. In this model, the depth of the trough would be related to the quantity of newly-produced metabolites from the peak that must be processed to restore metabolic balance. This is supported by the fact that oscillations did not result in overall more CO_2_ being fixed than the steady state (Table 1). Because oscillations are induced by following a period of no TPU limitation with induction of TPU limitation, it is possible that the plant has plentiful inorganic phosphate free during the start of the ramp. Then the excess is used to transiently surpass the TPU limitation of photosynthesis. The subversion of the steady state TPU-limited rate explains the similarity of ramps performed at 20°C and 25°C when the ramp speed is fast enough. TPU is the most temperature sensitive of the three components of photosynthesis (McClain & Sharkey, 2019), so a subversion of TPU limitation brings the photosynthetic rate back in line with the photosynthetic rate achieved at 25°C. Similarly, the plant should be able to dynamically exceed the RuBP regeneration limited portion of the curve if RuBP is initially in excess. The height of the peak would then be related to the size of the available metabolite pool.

The occurrence of oscillations suggests the existence of an “acute” TPU crisis that is rarely seen in the steady-state. Reduction of PSI without a corresponding increase in electron flow from the cytochrome *b6f* complex means that NADP^+^ must be limiting (Figure 6). This situation could occur if there is insufficient ATP production to process PGA into downstream products, limiting the flux through the reduction step. The troughs in assimilation are lower than the steady-state and are caused by a lack of ATP, caused by a sudden crisis in inorganic phosphate availability. This conclusion is supported by the decline in ATPase conductivity to protons (Figure 6). This acute restriction shows the photosynthetic rate as limited by a rapidly changing TPU limitation in response to phosphate levels, as opposed to the steady-state, which shows only the steady-state rate determined by the regulatory features that limit photosynthesis in response to TPU limitation. This data supports the conclusions of Laisk *et al*. (1991), who also found reduction of P_700_ during oscillations and calculated that NADPH/NADP^+^ ratios were antiparallel with oscillations in both photosynthesis and in ATP/ADP ratios. In the stroma, phosphate must be at a lowered concentration to maximize sucrose (Huber & Huber, 1996) and starch synthesis (Preiss, 1982), but must remain at a sufficient concentration to drive ATP synthesis. In acute TPU limitation, the balance is disrupted by a short period of very high photosynthetic rate. The transition from rubisco-limited to RuBP regeneration-limited conditions and vice versa involves much simpler adjustments in metabolism and so rarely produce oscillations.

The presence of an acute TPU crisis explains some non-obvious facets of steady-state TPU limitation. Triose phosphates do not necessarily build up in steady-state TPU limitation (Sharkey *et al*., 1986b), a counterintuitive fact considering it is the first output of a cycle that, according to the model, is going too fast. Instead, it is common that RuBP builds up, which is unexpected as TPU limitation implicitly limits the ATPase and RuBP requires ATP to be regenerated. The lack of ATP causes PGA to increase by as much as 77% and RuBP pools shrink immediately after the imposition of TPU but RuBP recovers as rubisco is deactivated (Sharkey *et al*., 1986b) and presumably other regulatory mechanisms are engaged. It will take additional studies of the effect of transients in metabolite pools to examine these regulatory mechanisms.

The amplitude of the oscillations is affected by several factors. The plants will not begin oscillating unless they enter TPU limitation suddenly. Ramps that are too slow allow time for complex adjustments in metabolism and so do not induce oscillations, and the amplitude of the oscillations varies with the speed at which the plants are induced into TPU limitation. This is emphasized in Figure 3, where the size of the overshoot varies with the length of time required to reach the beginning of oscillations. Plants ramped through an *A/C_i_* curve at low temperature are particularly susceptible and will oscillate with greater amplitude. The greatest amplitude is seen in the initial overshoot, and if the initial peak does not overshoot, there are no oscillations seen (for instance, the 100 ppm min^-1^ and 200 ppm min^-1^ ramps in Figure 3). If we believe that oscillations are caused by acute TPU limitation, the height of the overshoot will be related to the available metabolite pools usable before reaching a crisis in phosphate metabolism. When the ramp speed is fast, the integral of photosynthesis has been lower leading up to the beginning of oscillations, which would mean that the sum of metabolites consumed during the ramp is lower, while the potential to produce said metabolites should be approximately the same. When the plant reaches a *C_i_* that would typically cause RuBP regeneration or TPU limitation, greater pool sizes would produce a higher peak.

If TPU limitation in the steady state is best described as a collection of regulatory components (McClain & Sharkey, 2019), these oscillations are the result of the time delay to activate those regulatory components. The strength of the perturbation is important to the phenomenology because it puts strain on photosynthetic regulation. Oscillations are damped over a period of a few minutes, enough time to activate *PMF*-dependent control through energy-dependent quenching and photosynthetic control at cytochrome *b6f* (Kramer & Crofts, 1993, 1996), as well as rubisco deactivation which can begin in the first minutes of elevated CO_2_ (Sage *et al*., 1988) or just one min of exposure to low O_2_ to induce TPU (Sharkey *et al*., 1986b). Oscillations are seen when photosynthetic regulation is too slow to keep up with the changes in *A* and are damped when given enough time to activate regulatory controls on a timescale of min. This observation is supported by the reduced damping rate in oscillations induced via ramp (Table 2). The constantly changing setpoint for regulation causes the plant to perform worse and recover more slowly. This could explain the observation that rapid, large changes in instantaneous CO_2_ concentration in Free Air CO_2_ Enrichment studies may underestimate the improvement of photosynthesis and plant growth compared to conditions in which the CO_2_ does not change quickly (Allen *et al*., 2020).

## 5. Conclusions

TPU limitation shows flexibility during dynamic assimilation measurements, for precisely the same reason it is insensitive to O_2_ and CO_2_ changes: it is separated from rubisco by layers of metabolites. In the steady-state, inorganic phosphate pools are quite low (Sharkey & Vanderveer, 1989), but regulatory features balance the flux of inorganic phosphate into and out of the organic phosphate pool. Changing these fluxes dynamically imbalances photosynthesis and causes alternatively better and worse photosynthetic rate, and slower regulatory control is required to stabilize the photosynthetic rate again. This situation is a more intuitive understanding of TPU limitation – rather than being determined by a series of regulatory steps, the photosynthetic rate is determined by a crisis in metabolic pools.

At this point it may be useful to divide the phenomenon of TPU limitation into two separate categories. In the steady state, TPU-limited photosynthesis is described primarily by regulatory features such as rubisco deactivation and reduced electron flow because of energy-dependent quenching. In the acute, however, the photosynthetic rate temporarily defies some assumptions of the three-limitation model of steady-state photosynthesis. Dynamic TPU limitation must be controlled by pool sizes, and it is reflected in electron transport dynamics.

## Author contribution

AMM conceived the experiments and wrote the paper, TDS edited the paper and oversaw the research.

## Data Availability

All data are available from Dryad (address to be supplied later, deposit in process)

## Acknowledgements

Optical measurements reported here were enabled by instruments developed by David M. Kramer. Jeffrey A. Cruz and Robert Zegarac provided expertise on optical measurements.

